# BEVA2.0: Modular Assembly of Golden Gate-Compatible Vectors with Expanded Utility for Genetic Engineering

**DOI:** 10.1101/2024.12.04.626608

**Authors:** Barney A. Geddes, Riley Williamson, Jake Schumacher, Ahmad Ardi, Garrett Levin, Emily Červenka, Rui Huang, George C. diCenzo

## Abstract

This expansion for the modular vector assembly platform BEVA (Bacterial Expression Vector Archive) introduces 11 new BEVA parts including two new cloning site variants, two new antibiotic resistance modules, three new origins of replication, and four new accessary modules. As a result, the modular system is now doubled in size and expanded in its capacity to produce diverse replicating plasmids. Furthermore, it is now amenable to genetic engineering methods involving genome-manipulation of target strains through deletions or integrations. In addition to introducing the new modules, we provide several BEVA-derived Golden Gate cloning plasmids that are used to validate parts and that may be useful for genetic engineering of proteobacteria and other bacteria. We also introduce new parts to allow compatibility with the CIDAR MoClo parts libraries.

## Introduction

Since its inception, Golden Gate DNA assembly technology has revolutionized the ability to rapidly assemble multiple fragments of DNA in linear orders, facilitating advancements in synthetic biology (Engler *et al*., 2009). This technique, particularly through the MoClo (Modular Cloning) system, has enabled the hierarchical assembly of genetic constructs using standardized parts, making it a powerful tool for researchers. MoClo’s early adoption was largely driven by plant science research, where it was used to develop constructs for plant engineering via *Agrobacterium* tDNA integration (Werner *et al*., 2012). This led to the creation of extensive parts libraries and kits for plant engineering, such as those available through Addgene, which include promoters, untranslated regions, antigen tags, subcellular localization signals, marker genes, and terminators (Weber *et al*., 2011; Werner *et al*., 2012; Engler *et al*., 2014; Gantner *et al*., 2018), as well as tools for genome editing (Hahn *et al*., 2020; Grützner *et al*., 2021; Stuttmann *et al*., 2021). These resources have significantly accelerated synthetic biology by allowing researchers to rapidly assemble complex constructs using standardized parts.

While initially focused on plant engineering, the MoClo system naturally expanded to other organisms, including microorganisms. For example, parts kits have been developed for yeast hierarchical assembly (Lee *et al*., 2015) and protein expression (Obst, Lu and Sieber, 2017), and the CIDAR MoClo Library was developed for bacteria (Iverson *et al*., 2016), which includes a variety of useful synthetic biology parts sourced from the iGEM Registry of Standard Biological Parts. A CIDAR MoClo Expansion was also released that includes even more promoters, ribosome binding sites, coding sequences, and terminators, further enhancing its utility in synthetic biology. Useful systems for protein expression in *Escherichia coli* have also been developed (Moore *et al*., 2016; Bentham *et al*., 2021). However, with the widespread adoption of Golden Gate cloning, and new parts libraries in research labs around the world, there has been a growing need for new Golden Gate-compatible vectors with utility in a wide variety of organisms. To address this, we previously developed the Bacterial Expression Vector Archive (BEVA), a system that enables the assembly of Golden Gate cloning vectors from a library of standardized parts using a MoClo-like hierarchical assembly approach (Geddes, Mendoza-Suárez and Poole, 2019).

The field of microbial engineering continually seeks more efficient and versatile methods for designing plasmids to deliver and express genetic cargo. BEVA and the Standard European Vector Architecture (SEVA) (Silva-Rocha *et al*., 2013), which was recently adapted for Golden Gate cloning (Martínez-García *et al*., 2023), have emerged as promising tools due to their modular approach and ease of vector design, offering a path toward high-throughput applications. These models streamline the process of assembling complex genetic constructs by providing standardized, interchangeable modules that can be easily arranged in various configurations.

In this work, we report the creation of 11 new modules that significantly expand the utility of BEVA. We also generate and validate the performance of several new Golden Gate vectors that utilize the modules. The new vectors expand the utility of BEVA in two ways. First, they increase the capacity to generate useful expression vectors for diverse bacterial taxa by incorporating additional antibiotic resistance genes and broad host range origins of replication. Second, they incorporate modules that allow the generation of vectors useful for bacterial genome engineering, either through homologous or site-specific recombination. Overall, this study not only expands the BEVA toolkit but also contributes to the broader goal of developing versatile and efficient plasmid systems for microbial engineering.

## Materials and Methods

### Design of new BEVA modules

New BEVA modules were designed based on functional units from pre-existing cloning vectors and synthesized free of internal BsaI, BpiI (BbsI) and BsmBI (Esp3I) sites by Twist Bioscience in the Amp^R^ backbone “pTwist Amp High Copy”, Cm^R^ backbone “pTwist Chlor High Copy, or cloned via BsaI into Cm^R^ pGGAselect (New England Biolabs). In some cases, prototypes were designed by PCR amplification and cloning into pJet1.2, with the resulting vectors then used for generating Level 1 cloning vectors (pQGG002-005). However, these modules were later recloned into pGGAselect or resynthesized in the pTwist Chlor/Amp High Copy plasmids to eliminate a BsaI site from the pJet1.2 backbone that caused cloning efficiency issues.

#### Golden Gate cloning sites

Two new Level 1 cloning site modules were generated by altering the Level 1 cloning site from pOGG004. For pNDGG021 and pNDGG022, we excluded the T0 terminator 3’ of the cloning site. For pNDGG022, we further altered the 5’ BsmB1(Esp31) fusion site to CAGA to allow by-passing of the endlinker/terminator modules that include the *rrnB* T1 terminator and to allow for directly connecting these modules to a Position 4 module 3’ fusion site, thus completely excluding terminators flanking Level 1 cloning sites.

#### Antibiotic resistance cassettes

Antibiotic resistance modules were flanked by 5’ BsmB1(Esp31)-GCAA and 3’ ACTA-BsmB1(Esp31) sites to enable cloning as Position 2 modules in BEVA. For the kanamycin/neomycin resistance gene *nptII* (in pNDGG001), we used 1,036 bp from the cloning vector pTH1937 (Milunovic *et al*., 2014) that included the *nptII* open reading frame and 241 bp upstream which was cloned into pGGAselect via BsaI. For the spectinomycin resistance gene *aadA* (in pNDGG002), we synthesized 1,786 bp from the pHP45Ωspec interposon (Prentki and Krisch, 1984) that contained the *aadA* open reading frame with 516 upstream and 259 bp downstream regions in pTwist Amp High Copy.

#### Origins of replication and transfer

Three new origins of replication and transfer were generated. These modules were each flanked by 5’ BsmB1(Esp31)-ACTA and 3’ TTAC-BsmB1(Esp31) sites to enable cloning as Position 3 modules in BEVA. The sequences of the origins p15A (pNDGG006) and pMB1 (pNDGG007), including their *oriT*s, were derived from 1,492 bp and 980 bp segments from pTH1937 (Milunovic *et al*., 2014) and pUCP30T (Schweizer and Hoang, 1996), respectively. pMB1 was cloned into pGGAselect via BsaI, while p15A was synthesized in pTwist Chlor High Copy. The broad host range origin RSF1010 (3,078 bp) was PCR amplified from pRSF-LtetO-GFP (Lee *et al*., 2019) and combined with a 268 bp *oriT* amplified from pOGG026 (Geddes, Mendoza-Suárez and Poole, 2019) by BsaI Golden Gate cloning into pGGAselect to generate pNDGG005.

#### Accessory modules

Four accessory modules were added, each flanked with 5’ BsmB1(Esp31)-TTAC and 3’ CAGA-BsmB1(Esp31) sites to enable cloning as Position 4 modules in BEVA. The *sacB* gene enabling sucrose counterselection was designed based off of pJQ200SK (Quandt and Hynes, 1993), and the resulting module in pNDGG008 included 1,944 bp surrounding the *sacB* open reading frame including 446 bp upstream and 76 bp downstream.

The *FRT* site-specific Flp recombinase motif (GAAGTTCCTATTCCGAAGTTCCTATTCTCTAGAAAGTATAGGAACTTC) was synthesized as a Position 4 module in pNDGG009 with a few extra flanking base pairs to bring the total size to 63 bp. The *attP* φC31 integrase target site (CCCCAACTGGGGTAACCTTTGAGTTCTCTCAGTTGGGGG) was similarly synthesized as a Position 4 module in pNDGG010. An *attP* motif was also included in pNDGG009 flanked by BsaI (5’GCTT and 3’CGCT) sites (CIDAR EF module) such that it could be added to the 3’ end of an open reading frame in a Level 1 cloning. Likewise, a *FRT* motif flanked by the same BsaI sites (5’GCTT/3’CGCT) was added to pNDGG010.

The I-SceI site accessary module in pQGG001 was generated by ligating the I-SceI site as an annealed double-stranded oligomer (5’ – TCTGGACTACGGTTCCAAATTACCCTGTTATCCCTACCTTGGAATGGTCA-3’ and 5’-TTACTGACCATTCCAAGGTAGGGATAACAGGGTAATTTGGAACCGTAGTC-3’) into an *Esp3I*-digested pOGG012 backbone using T4 DNA ligase.

#### CIDAR terminators

Three terminators from CIDAR (T2m, T12m and T15m from CIDAR MoClo Extension Volume I) were adapted such that the 3’ fusion site matched BEVA L1 cloning vectors. The fusion site was adapted during cloning as a Level 0 part into pOGG006. The same forward primer was used for all three, which bound upstream of the 5’ BsaI site and added a BpiI fusion site (TGCC) for cloning into pOGG006 (5’-AAAGAAGACAATGCCCGGCCGCTTCTAGAGA-3’). A unique 3’ primer was used for each, with the binding site anchored in the terminator and an altered 3’ BsaI fusion site added in the primer along with a BpiI fusion site for cloning into pOGG006 (T2m = 5’-AAAGAAGACAATCCCGGTCTCAAGCGTCTCAAGGGCGCAATAAAA-3’; T12m = 5’-AAAGAAGACAATCCCGGTCTCAAGCGTTGAGAAGAGAAAAGAAAACCG- 3’, T15m = 5’-AAAGAAGACAATCCCGGTCTCAAGCGGGCAGACCAGAAACAAA-3’. After cloning into pOGG006, the resulting plasmids functioned as Level 0 parts, for BEVA, flanked by *BsaI* and the fusion sites GCTT (5’) and CGCT (3’).

### Golden Gate cloning

Golden Gate cloning reactions were performed using forty fmol of plasmid parts in a reaction mix recipe of 1 µl restriction enzyme (BsmBI-v2, Esp3I, or BsaI-HF-v2) (New England Biolabs), 1.5 µl bovine serum albumin, 1 µl T4 DNA ligase (400 units/µL; New England Biolabs), and 1.5 µl 10x T4 DNA ligase buffer, diluted with ddH_2_O to a total volume of 15 µl. Alternatively, NEB Golden Gate Assembly Kits (BsaI-HF-v2 or BsmBI-v2) were used as a pre-mixed reaction mix with the DNA parts and ddH_2_O. The Golden Gate cloning reactions were performed in a thermocycler as follows: For BbsI/Esp3I: 30 cycles of 42°C for 1 minute followed by 16°C for 1 minute, with heat inactivation at 60°C for 5 minutes; for BsaI: 30 cycles of 37°C for 1 minutes and 16°C for 1 minute, with heat inactivation at 60°C for 5 minutes. The resulting reaction mixes were stored at-20°C before thawing and using to transform chemically competent *E. coli* DH5α. Successful transformants were selected using appropriate antibiotics and X-gal (5-bromo-4-chloro-3-indolyl-β-D-galactopyranoside). Successful vector constructions were selected based on blue-colored colonies, while successful Level 1 clonings were selected based on white-colored colonies. Plasmids were verified by diagnostic restriction digest and/or whole plasmid long-read sequencing by Plasmidsaurus.

### Genetic manipulations

Bacterial strains used in this work are listed in Table 1. *E. coli* and *Sinorhizobium meliloti* strains were grown in liquid or solid Lysogeny Broth (LB) medium, supplemented with 0.25 mM MgCl2₂ and 0.25 mM CaCl₂ (termed LBmc) for *S. meliloti. E. coli* strains were grown at 37°C while *S. meliloti* strains were grown at 28°C. Antibiotics were added to media when necessary for *E coli* (Ec) or *S. meliloti* (Sm), at the following concentrations: gentamicin 10 μg/ml (Ec) or 60 μg/ml (Sm), kanamycin 25 μg/ml (Ec), neomycin 200 μg/ml (Sm), spectinomycin 50 μg/ml (Ec) or 100 μg/ml (Sm), ampicillin 100 μg/ml (Ec), and tetracycline 5 μg/ml (Ec or Sm). Sucrose counterselection was performed by adding 10% sucrose to LB medium. Conjugations between *E. coli* and *S. meliloti* were performed by triparental mating using *E. coli* MT616 as a helper strain (Finan *et al*., 1986). Tests for colony antibiotic resistance/sensitivity phenotypes were performed by patching colonies onto replicated agar plates using a sterilized toothpick. Transformation into *E. coli* and conjugation into *S. meliloti* by triparental mating were performed using routine protocols (Finan *et al*., 1986; Geddes, Mendoza-Suárez and Poole, 2019).

**Table 1.** Plasmids used in this work.

| Vector Name | Description | Citation |
| --- | --- | --- |
| *Vector construction parts used to assemble BEVA2.0 vectors* |  |  |
| pOGG004 | BEVA Position 1 Level 1 BsaI cloning site flanked by T0, Sp<sup>R</sup>. Addgene 113979 | (Geddes, Mendoza-Suárez and Poole, 2019) |
| pOGG006 | BEVA Position 1 Level 2 BpiI cloning site flanked by T0, Sp<sup>R</sup>. Addgene 113981 | (Geddes, Mendoza-Suárez and Poole, 2019) |
| pNDGG021 | BEVA2.0 Position 1 Level 1 BsaI cloning site without T0, Amp<sup>R</sup> | This work |
| pNDGG022 | BEVA2.0 Position 1 Level 1 BsaI cloning site with alternate CAGA 3' fusion site, without T0, Amp<sup>R</sup> | This work |
| pOGG008 | BEVA Position 2 Km/Nm antibiotic resistance via *aphA*, Sp<sup>R</sup>/Km<sup>R</sup>/Nm<sup>R</sup>. Addgene 113982 | (Geddes, Mendoza-Suárez and Poole, 2019) |
| pOGG009 | BEVA Position 2 Gm antibiotic resistance via *aac3*, Sp<sup>R</sup>/Gm<sup>R</sup>. Addgene 113983 | (Geddes, Mendoza-Suárez and Poole, 2019) |
| pOGG042 | BEVA Position 2 Gm antibiotic resistance via *aac3*, Sp<sup>R</sup>/Gm<sup>R</sup>. Addgene 113998 | (Geddes, Mendoza-Suárez and Poole, 2019) |
| pNDGG001 | BEVA2.0 Position 2 Km/Nm antibiotic resistance via *nptII*, Cm<sup>R</sup>/Km<sup>R</sup>/Nm<sup>R</sup>. | This work |
| pNDGG002 | BEVA2.0 Position 2 Sp antibiotic resistance via *aadA*, Amp<sup>R</sup>/Sp<sup>R</sup> | This work |
| pOGG010 | BEVA Position 3 RK2 broad host range origin of replication with *oriT*, Sp<sup>R</sup>. Addgene 113984 | (Geddes, Mendoza-Suárez and Poole, 2019) |
| pOGG011 | BEVA Position 3 pBBR1 broad host range origin of replication with *oriT*, Sp<sup>R</sup>. Addgene 113985 | (Geddes, Mendoza-Suárez and Poole, 2019) |
| pNDGG005 | BEVA2.0 Position 3 RSF1010 broad host range origin of replication with *oriT*, Cm<sup>R</sup>. | This work |
| pNDGG006 | BEVA2.0 Position 3 p15A narrow host range origin of replication with *oriT*, Cm<sup>R</sup>. | This work |
| pNDGG007 | BEVA2.0 Position 3 pMB1 narrow broad host range origin of replication with *oriT*, Cm<sup>R</sup>. | This work |
| pOGG012 | BEVA Position 4 *parCDABE* stability locus, Sp<sup>R</sup>. Addgene 113986 | (Geddes, Mendoza-Suárez and Poole, 2019) |
| pNDGG008 | BEVA2.0 Position 4 *sacB* sucrose counterselection module, Amp<sup>R</sup>. | This work |
| pQGG001 | BEVA2.0 Position 4 I-SceI recognition/cut site, Sp<sup>R</sup>. | This work |
| pNDGG009 | BEVA2.0 Position 4 *FRT* recombinase site, Amp<sup>R</sup>. Also includes *attP* EF module for L1 GG cloning. | This work |
| pNDGG010 | BEVA2.0 Position 4 *attP* recombinase site, Amp<sup>R</sup>. Also includes *FRT* EF module for L1 GG cloning. | This work |
| pOGG013 | BEVA Position 3 endlinker with T1 terminator, Sp<sup>R</sup>. Addgene 113987 | (Geddes, Mendoza-Suárez and Poole, 2019) |
| pOGG014 | BEVA Position 4 endlinker with T1 terminator, Sp<sup>R</sup>. Addgene 113988 | (Geddes, Mendoza-Suárez and Poole, 2019) |
| pOGG015 | BEVA Position 5 endlinker with T1 terminator, Sp<sup>R</sup>. Addgene 113989 | (Geddes, Mendoza-Suárez and Poole, 2019) |
| pOGG016 | BEVA Position 6 endlinker with T1 terminator, Sp<sup>R</sup>, Addgene 113990 | (Geddes, Mendoza-Suárez and Poole, 2019) |
| ***BEVA2.0 Vectors used for Level 1 or 2 GG cloning*** |  |  |
| pNDGG003 | BEVA2.0 Broad host range plasmid with stability. L1-RK2-Tc-par-ELT4; GG assembly from pOGG010, pOGG012, pOGG014, pOGG042, and pOGG004, Tc<sup>R</sup> | This work |
| pNDGG004 | BEVA2.0 Broad host range plasmid with stability. L1-pBBR1-Tc-par-ELT4; GG assembly from pOGG011, pOGG012, pOGG014, pOGG042, and pOGG004, Tc<sup>R</sup> | This work |
| pNDGG049 | BEVA2.0 Broad host range plasmid with stability. -pBBR1-Gm-par-ELT4; GG assembly from pOGG004, pOGG009, pOGG011, pOGG012, and pOGG014, Gm<sup>R</sup> | This work |
| pNDGG050 | BEVA2.0 Broad host range plasmid with stability. L1-RK2-Gm-par-ELT4; GG assembly from pOGG004, pOGG009, pOGG010, pOGG012, and pOGG014, Gm<sup>R</sup> | This work |
| pNDGG051 | BEVA2.0 Broad host range plasmid with stability. L1-pBBR1-Sp-par-ELT4; GG assembly from pOGG004, pNDGG002, pOGG011, pOGG012, and pOGG014, Sp<sup>R</sup> | This work |
| pNDGG052 | BEVA2.0 Broad host range plasmid with stability. L1-RK2-Sp-par-ELT4; GG assembly from pOGG004, pNDGG002, pOGG010, pOGG012, and pOGG014, Sp<sup>R</sup> | This work |
| pNDGG053 | BEVA2.0 Broad host range plasmid with stability. L1-pBBR1-Km-par-ELT4; GG assembly from pOGG004, pNDGG001, pOGG011, pOGG012, and pOGG014, Nm<sup>R</sup>/Km<sup>R</sup> | This work |
| pNDGG054 | BEVA2.0 Broad host range plasmid with stability. L1-RK2-Km-par-ELT4; GG assembly from pOGG004, pNDGG001, pOGG010, pOGG012, and pOGG014, Nm<sup>R</sup>/Km<sup>R</sup> | This work |
| pNDGG056 | BEVA2.0 Broad host range plasmid with stability. L1-RSF1010-Gm-par-ELT4; GG assembly from pOGG004, pOGG009, pNDGG005, pOGG012, and pOGG014, Gm<sup>R</sup> | This work |
| pNDGG057 | BEVA2.0 Broad host range plasmid with stability. L1-RSF1010-Tc-par-ELT4; GG assembly from pOGG004, pOGG042, pNDGG005, pOGG012, and pOGG014, Tc<sup>R</sup> | This work |
| pNDGG058 | BEVA2.0 Broad host range plasmid with stability. L1-RSF1010-Sp-par-ELT4; GG assembly from pOGG004, pNDGG002, pNDGG005, pOGG012, and pOGG014, Sp<sup>R</sup> | This work |
| pNDGG059 | BEVA2.0 Broad host range plasmid with stability. L1-RSF1010-Km-par-ELT4; GG assembly from pOGG004, pNDGG001, pNDGG005, pOGG012, and pOGG014, Nm<sup>R</sup>/Km<sup>R</sup> | This work |
| pNDGG070 | BEVA2.0 Sucrose curable broad host range plasmid . L1-pBBR1-Sp-sacB-ELT4; GG assembly from pOGG004, pNDGG002, pOGG011, pNDGG008, and pOGG014, Sp<sup>R</sup> | This work |
| pQGG002 | BEVA2.0 Narrow host range plasmid with I-SceI counterselectable site. L1-P15A-Nm-ISceI; GG assembly from pOGG004, pNDGG001, pNDGG006, pQGG001, pOGG014. Km<sup>R</sup>/Nm<sup>R</sup>. | This work |
| pQGG003 | BEVA2.0 Narrow host range plasmid with I-SceI counterselectable site for BpiI cloning. L2-P15A-Nm-ISceI; GG assembly from pOGG006, pNDGG001, pNDGG006, pQGG001, pOGG014. Km<sup>R</sup>Nm<sup>R</sup>. | This work |
| pQGG004 | BEVA2.0 Narrow host range plasmid with I-SceI counterselectable site. L1-P15A-Gm-ISceI-ELT4; GG assembly from pOGG004, pOGG009, pNDGG006, pQGG001, pOGG014. Gm<sup>R</sup>. | This work |
| pQGG005 | BEVA2.0 Narrow host range plasmid with I-SceI-ELT4 counterselectable site for BpiI cloning. L2-P15A-Gm-ISceI-ELT4; GG assembly from pOGG006, pOGG009, pNDGG006, pQGG001, pOGG014. Gm<sup>R</sup>. | This work |
| pNDGG012 | Narrow host range plasmid with *FRT* site. L1-pMB1-Gm-*FRT*-ELT4; GG assembly from pNDGG021, pOGG009, pNDGG007, pNDGG009, pOGG014. Gm<sup>R</sup>. | This work |
| pNDGG013 | Narrow host range plasmid with *FRT* site. L1-p15A-Km-*FRT*-ELT4; GG assembly from pNDGG021, pOGG009, pNDGG006, pNDGG009, pOGG014. Km<sup>R</sup>. | This work |
| pNDGG014 | Narrow host range plasmid. L1-pMB1-Gm-ELT3; pNDGG021, pOGG009, pNDGG007, pOGG013. Gm<sup>R</sup>. | This work |
| pNDGG015 | Narrow host range plasmid. L1-p15A-Km-ELT3; pNDGG021, pOGG009, pNDGG006, pOGG013. Km<sup>R</sup>. | This work |
| **ORF parts used for BsaI GG cloning** |  |  |
| J23106 | CIDAR Medium-strength constitutive promoter from Anderson series, AB module, Amp<sup>R</sup>, Addgene 65992 | (Iverson *et al.*, 2016) |
| B0032m | CIDAR Weiss RBS of medium strength, BC module, Amp<sup>R</sup>, Addgene 66020 | (Iverson *et al.*, 2016) |
| BCD12 | CIDAR BiCistronic RBS of medium-low strength, BC module, Amp<sup>R</sup>, Addgene 66023 | (Iverson *et al.*, 2016) |
| C3m | CIDAR MoClo Extension volume I, sfGFP, CD module, Amp<sup>R</sup>. Addgene 120956 | Gift from Richard Murray via Addgene) |
| C51m | CIDAR MoClo Extension volume I, sfYFP, CD module, Amp<sup>R</sup>. Addgene 120975 | Gift from Richard Murray via Addgene) |
| C91m | CIDAR MoClo Extension volume I, sfCFP, CD module, Amp<sup>R</sup>. Addgene 120999 | Gift from Richard Murray via Addgene) |
| C88m | CIDAR MoClo Extension volume I, m-Scarlet-I, CD module, Amp<sup>R</sup>. Addgene 121003 | Gift from Richard Murray via Addgene) |
| T2m | CIDAR MoClo Extension volume I, Eck120033736 terminator (nmeth.2515), DE module, Amp<sup>R</sup>. Addgene 121029 | Gift from Richard Murray via Addgene) |
| T12m | CIDAR MoClo Extension volume I, Eck120026300 terminator (nmeth.2515), DE module, Amp<sup>R</sup>. Addgene 121031 | Gift from Richard Murray via Addgene) |
| T15m | CIDAR MoClo Extension volume I, L3S3P11 terminator (nmeth.2515), DE module, Amp<sup>R</sup>. Addgene 121034 | Gift from Richard Murray via Addgene) |
| pOGG037 | BEVA1.0 sfGFP “PU” model (CE CIDAR extensions), Sp<sup>R</sup>. Addgene 113995 | (Geddes, Mendoza-Suárez and Poole, 2019) |
| pNDGG037 | BEVA2.0 T2m terminator with DF extension for CIDAR MoClo Golden Gate Assembly, DF module. Cloned via BpiI GG Assembly of PCR amplicon into POGG006, Sp<sup>R</sup>. | This work |
| pNDGG038 | BEVA2.0 T12m terminator with DF extension for CIDAR MoClo Golden Gate Assembly, DF module. Cloned via *BpiI* GG Assembly of PCR amplicon into POGG006, Sp<sup>R</sup>. | This work |
| pNDGG039 | BEVA2.0 T15m terminator with DF extension for CIDAR MoClo Golden Gate Assembly, DF module. Cloned via *BpiI* GG Assembly of PCR amplicon into POGG006, Sp<sup>R</sup>. | This work |
| **Other plasmids** |  |  |
| pDA1-SceI-sacB | pBBR1 broad host range plasmid expressing I-SceI homing endonuclease and *sacB* counterselectable marker, Tc<sup>R</sup> | (Aubert, Hamad and Valvano, 2014) |
| pNDMS450 | RK2 broad host range plasmid expressing constitutive sfGFP. GG assembly of pNDGG054_AF, J23106_AB, B0032m_BC, pOGG037_CE, and EF oligo, Tc<sup>R</sup> | This work |
| pNDMS451 | pBBR1 broad host range plasmid expressing constitutive sfGFP. GG assembly of pNDGG053_AF, J23106_AB, B0032m_BC, pOGG037_CE, and EF oligo, Tc<sup>R</sup> | This work |
| pNDMS459 | RSF1010 broad host range plasmid expressing constitutive sfGFP. GG assembly of pNDGG054_AF, J23106_AB, B0032m_BC, pOGG037_CE, and EF oligo, Tc<sup>R</sup> | This work |
| pNDMS054 | RK2 broad host range plasmid expressing constitutive sfGFP. GG assembly from pNDGG003_AF, J23106_AB, BCD12_BC, C3m_CD, and T2m_DF (pNDGG037), Tc<sup>R</sup> | This work |
| pNDMS055 | RK2 broad host range plasmid expressing constitutive sfCFP. GG assembly from pNDGG003_AF, J23106_AB, BCD12_BC, C91m_CD, and T2m_DF (pNDGG037), Tc<sup>R</sup> | This work |
| pNDMS056 | RK2 broad host range plasmid expressing constitutive sfYFP. GG assembly from pNDGG003, J23106_AB, BCD12_BC, C51m_CD, and T2m_DF (pNDGG037), Tc<sup>R</sup> | This work |
| pNDMS057 | RK2 broad host range plasmid expressing constitutive m-Scarlet-I. GG assembly from pNDGG003, J23106_AB, BCD12_BC, C99m_CD, and T2m_DF (pNDGG037), Tc<sup>R</sup> | This work |

### Fluorescence measurements by flow cytometry and plate reader assays

Preparation of cells for flow cytometry experiments were conducted as follows. Cells were grown in LB media containing standard working concentrations of the antibiotic corresponding to the plasmid-encoded resistance. Media was inoculated with a single colony and placed on a shake incubator set to 37°C at 215 rpm and grown overnight. One ml of each cell culture was transferred to individual 1.5 ml microtubes, after which their optical density was measured using a spectrophotometer. The tubes were then centrifuged for one minute at 17,000 x g. The supernatant was removed, and the pellet was resuspended in PBS (phosphate-buffered saline) to a final OD600 (optical density at 600 nm) of 1.0. Resuspended cells were further diluted 1:100 in PBS immediately prior to flow cytometry. Tubes containing the diluted samples were placed in the flow cytometer receptacle one at a time. The Beckman Cytoflex S flow cytometer was set to interrogate the cells using yellow and blue lights. The yellow light detectors were set to measure an mScarlet-I band path of 585/42, and the blue light detectors were set to measure both GFP at 525/40 and SSC at 488/8. Speed settings were adjusted to reach an average of 1000 events per second. These settings were adjusted depending on each sample but were generally kept at the lowest speed of 10 µl/min. Samples were manually compensated when required.

For measurements of fluorescence by plate reader, a Cytation 5 was used to quantify fluorescence across a bacterial growth curve. *E. coli* or *S. meliloti* were grown at 37°C and 28°C, respectively, for 24 hours from a starting OD600 of 0.05. The OD600 and fluorescence was measured every 20 minutes. The growth was analyzed using the Growthcurver package in R. The fluorescent values presented are derived from the maximum doubling time in the growth curve. The following parameters were used for fluorescent measurements: cfCFP - excitation 485, emission 475, gain 90; sfGFP - excitation 485, emission 510, gain 60; sfYFP - excitation 510, emission 535, gain 100; mScarlet-I - excitation 569, emission 593, gain 108.

## Results and Discussion

### Expansion of BEVA with 11 new modules

We aimed to improve the utility of BEVA for the construction of useful Golden Gate vectors by (1) expanding the potential host range with additional broad host range origin of replication and antibiotic resistance modules, and (2) expand the utility to routinely used vectors for genome manipulations (deletions/integrations) via the addition of narrow host range origins of replication, counters-electable markers, and site-specific recombinase sites. To enable this, we designed 11 new modules that are shown as part of the broader BEVA archive in **Figure 1** and listed in **Table 1**. The rational for inclusion of each of the modules is described below.

**Figure 1.**
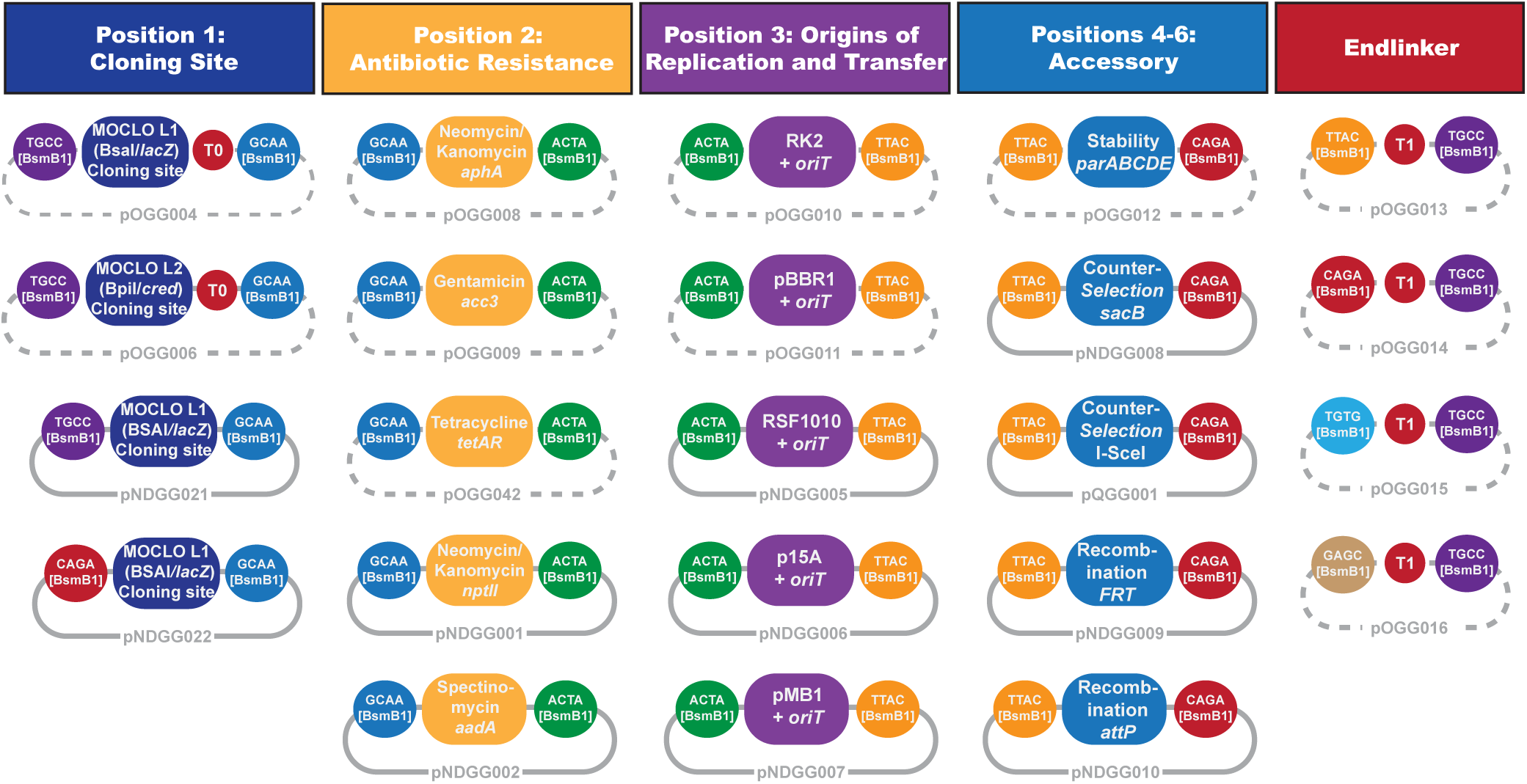
Diagram of vector modules in the expanded BEVA2.0 vector archive. Dashed vector backbones reflect modules from BEVA1.0 (Geddes, Mendoza-Suárez and Poole, 2019). Solid vector backbones reflect BEVA2.0 expansion modules presented in this work. Colored rounded rectangles reflect parts, colored by module type. Colored circles reflect sticky overhangs generated by *BsmB1* and ligated by DNA ligase in Golden Gate cloning reactions.

The initial BEVA was largely designed as a platform for generating expression vectors and hence included the broad host range origins of replication pBBR1 and RK2 with several options for antibiotic resistance genes that could be tailored to a target strain of interest. To facilitate wider host utility, we added another widely-used broad host range origin of replication, RSF1010 (vector pNDGG005), as a Position 3 module. We also expanded available antibiotic selection by adding spectinomycin resistance via *aadA* as a Position 2 module (vector pNDGG002). While a kanamycin/neomycin resistance module (*aphA*-based) was available in the original BEVA, we and others have found this module to be ineffective for neomycin-based selection in *Sinorhizobium meliloti* (personal communication). Therefore, a new kanamycin/neomycin module was added based on the more widely used *nptII* gene (vector pNDGG001).

Genome manipulations in model bacteria such as *S. meliloti* 1021 routinely involve the use of integrative, or “suicide”, plasmids that replicate readily in *E. coli* but must integrate into the genome to be maintained in the target microbe (Quandt and Hynes, 1993; diCenzo *et al*., 2019). Integrations can be catalyzed by cloning a short sequence of the target genome to facilitate integration at a specific genomic locus via homologous recombination. Double homologous recombination can generate deletions in a similar approach by including target-genome sites flanking a region to be deleted and incorporating a counter-selectable marker in the vector backbone to catalyze its excision. An alternative to homologous recombination is to utilize site-specific recombinases to catalyze integration at a target site in the host genome with a compatible recombinase target sequence of an integrative plasmid (diCenzo and Finan, 2018).

These approaches can also be combined to catalyze inversions or large deletions between two recombinase sites, following their integration on either side of a target region via two heterologous integrative vector backbones. To facilitate all of these approaches, we added two narrow host range origins (p15A and pMB1, vectors pNDGG006-7), two site-specific recombinase sites (*FRT* and *attP*, vectors pNDGG009-10), and two counter-selectable markers (I-SceI and *sacB*, vectors pQGG001 and pNDGG008, respectively). The I-SceI target sequence can be counter-selected through expression of the I*-*SceI restriction enzyme from an introduced plasmid (Aubert, Hamad and Valvano, 2014), while *sacB* can be counter-selected via sucrose toxicity (Quandt and Hynes, 1993). The *FRT* and *attP* sites facilitate recombination with *FRT* and *attP* sites via Flp recombinase and φC31 integrase, respectively (diCenzo and Finan, 2018). The availability of two heterologous narrow host range origins facilitates introductions of multiple plasmids by homologous recombination (such as for *FRT*-mediated deletions), minimizing the chance for recombination between them (diCenzo and Finan, 2018). To allow further minimization of homology between assembled plasmids, we also introduced three alternate Level 1 cloning sites (pNDGG021-23) that allow exclusion of the 3’ T0 terminator or the 5’ T1 terminator (via skipping an endlinker) from assembled vectors.

### A suite of broad host range Golden Gate expression vectors

Using the updated BEVA2.0 parts library, we combined several modules with the aim of developing a flexible broad host range vector suite with varied antibiotic resistance. To this end, we included four antibiotic resistance modules (*aadA* (Sp), *nptII* (Nm/Km), *tetAR* (Tc), and *acc3* (Gm); Position 2) with three broad host range origins of replication (RK2, pBBR1 and RSF1010; Position 3). These were combined in the standard BEVA architecture with a Level 1 Golden Gate cloning site (Position 1), the *par* accessary cluster for stability in the absence of antibiotic selection (Position 4), and an endlinker (ELT4). The resulting vectors and their compositions are summarized in **Table 1** and **Figure 2A**.

**Figure 2.**
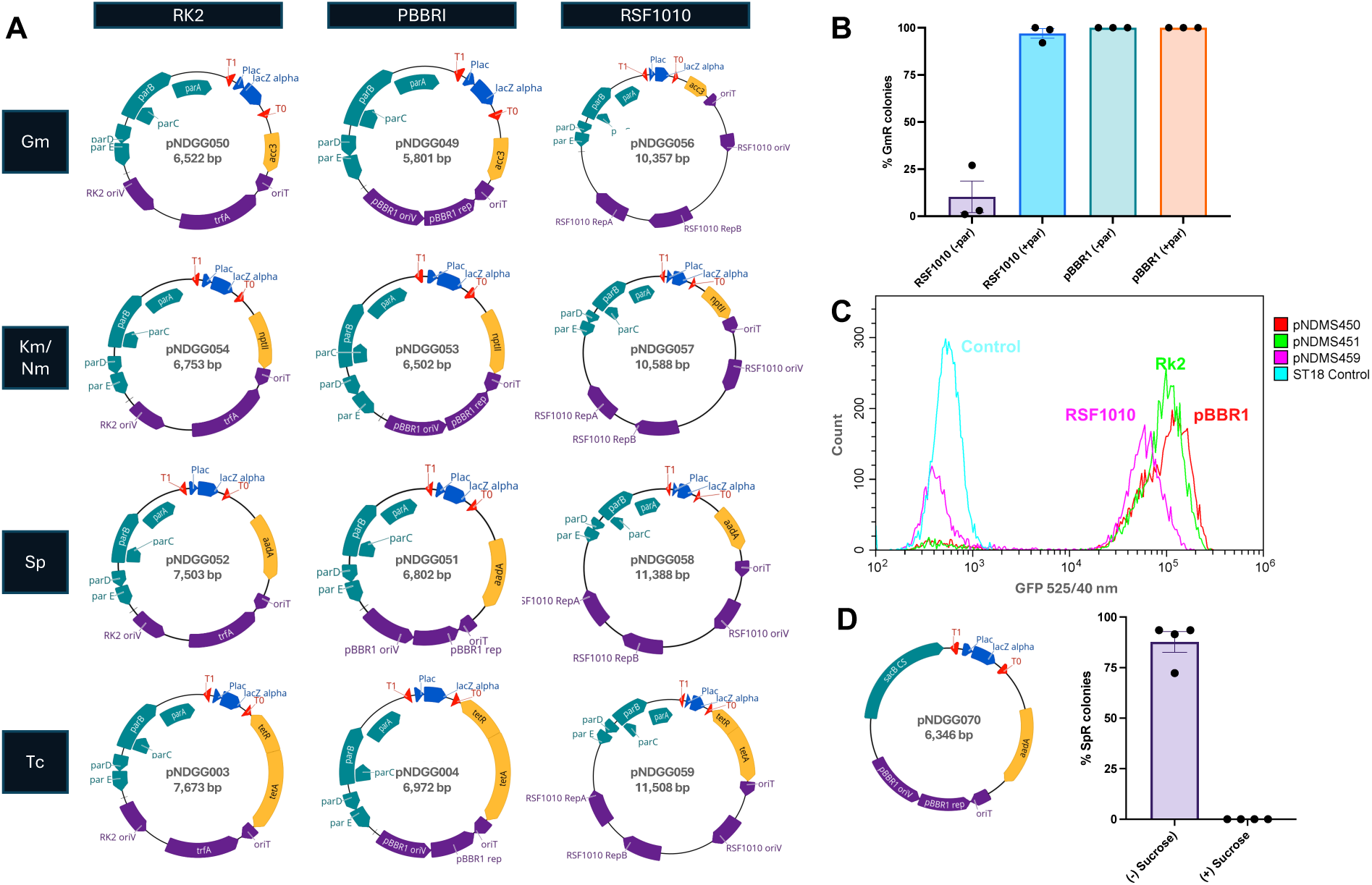
BEVA2.0 Broad Host Range Vectors. **A.** Plasmid maps of broad host range vectors developed using BEVA2.0 parts. Nucleotide sequences of the plasmids are included in the supplemental data. **B.** Quantification of pBBR1 or RSF1010 plasmid maintenance with and without *par* loci. Data is expressed as the proportion of colonies that grew on LB + Gm plates after patching following antibiotic-free culturing and spread-plating on LB. **C.** Histograms of GFP signal from flow cytometry analysis of *E. coli* ST18ALA bearing RSF1010, pBBR1, or RK2 plasmids expressing sfGFP. **D.** Plasmid map of a curable *sacB*-containing pBBR1 plasmid pNDGG070, and the proportion of plasmid loss following growth on 10% sucrose based on sensitivity when colonies were patched on LB + Sp media.

While the *parABCDE* cluster has been characterized for its ability to stabilize RK2 plasmids from which it was derived, we wished to verify the compatibility of this module with the heterologous origins of replication pBBR1 and RSF1010. To do so, we performed an experiment to assess plasmid stability in the absence of antibiotic selection, and compared the stability of plasmids that included the module (Lv1-Gm-pBBR1-par-ELT4, and Lv1-Gm- RSF1010-par-ELT4) to vectors we constructed the excluded the *par* module (Lv1-Gm-pBBR1- ELT3, and Lv1-Gm-RSF1010-ELT3). Single colonies of *E. coli* transformants from each plasmid were used to inoculate an overnight culture in the absence of antibiotics. Following overnight growth and saturation, the culture was plated by serial dilution to single colonies without antibiotic selection, and the resulting colonies were patched onto Gm to assess plasmid maintenance. In this experiment, while the pBBR1 vector showed stability in either the presence or absence of *par*, the RSF1010 plasmid was significantly more stable when including the *par* module (**Figure 2B**).

To verify Level 1 cloning and suitability of plasmids for expression, one plasmid from each origin of replication from the suite (all Tc^R^) were used for Level 1 Golden Gate cloning, wherein we combined sfGFP with a constitutive promoter and ribosome binding site in *E. coli* ST18ALA (**Table 3**). The fluorescence of the resulting plasmids was tested using flow cytometry. We found similar fluorescence profiles in plasmids that contained RK2, pBBR1, or RSF1010 backbones (**Figure 2C**).

We also demonstrated the flexibility of the BEVA system by cloning an alternate broad host range plasmid that included a module to facilitate its removal rather than promote stability, *sacB* (Figure 2B). We used a plasmid (Lv1-Sp-pBBR1-sacB-ELT4) to facilitate curing (i.e., loss) of the difficult to remove pBBR1 plasmids from strains of a *S. meliloti* deletion library (Milunovic *et al*., 2014) through incompatibility. Thereafter, we found that the BEVA2.0 *sacB* plasmid was efficiently removed by sucrose counterselection (**Figure 2E**). In four replicates, *S. meliloti* bearing the Sp^R^ *sacB* plasmid were plated onto LBmc or LBmc with 10% sucrose by serial dilution following overnight culturing in the absence of antibiotics. Approximately 50 single colonies from each set of plates were patched onto LBmc Sp to test for plasmid loss. A low rate of plasmid loss was observed in the absence of sucrose selection, while sucrose selection resulted in a complete curing of the plasmid in the population tested.

### Golden-Gate compatible vectors for homologous recombination

We also expanded the collection of Level 1 cloning vectors for use in genetic engineering by utilizing integrative plasmid parts from the updated BEVA2.0 parts library (Table 1). Vectors for homologous recombination were constructed using the p15A origin of replication, which generates integrative plasmids in many Gram-negative bacteria due to its inability to be maintained as a replicating plasmid (Quandt and Hynes, 1993) (Figure 3A). To facilitate workflows for constructing deletions by double homologous recombination, we also added an I*-* SceI recognition site that enables curing through expression of the I*-SceI* endonuclease. We generated multiple versions including optional antibiotic selection (Gm^R^ or Nm^R^), and two different cloning sites (BsaI or BpiI) to facilitate flexible cloning of chromosomal homologous regions, should one site or the other be present in a target region. These plasmids have been used in workflows to generate deletions in *S. meliloti* and functioned efficiently to generate marked or unmarked deletions. For example, when a pQGG004 derivative was used to create a marked deletion of the *S. meliloti phaZ* gene, the efficiency of I*-*SceI selection for the expected double-recombination was ∼14% when averaged across four independent replicates with 17-33 colonies. Surprisingly, the efficiency varied from as low as 3% to up to 19% depending on the trial, which is a result that we have repeatedly observed when selecting for double recombinants using this approach (diCenzo, unpublished). We hypothesize that the efficiency of obtaining double recombinants varies depending on where the initial recombination occurred, and thus we recommend purifying multiple single recombinants for use in downstream steps to optimize the likelihood of obtaining a correct double recombinant.

**Figure 3.**
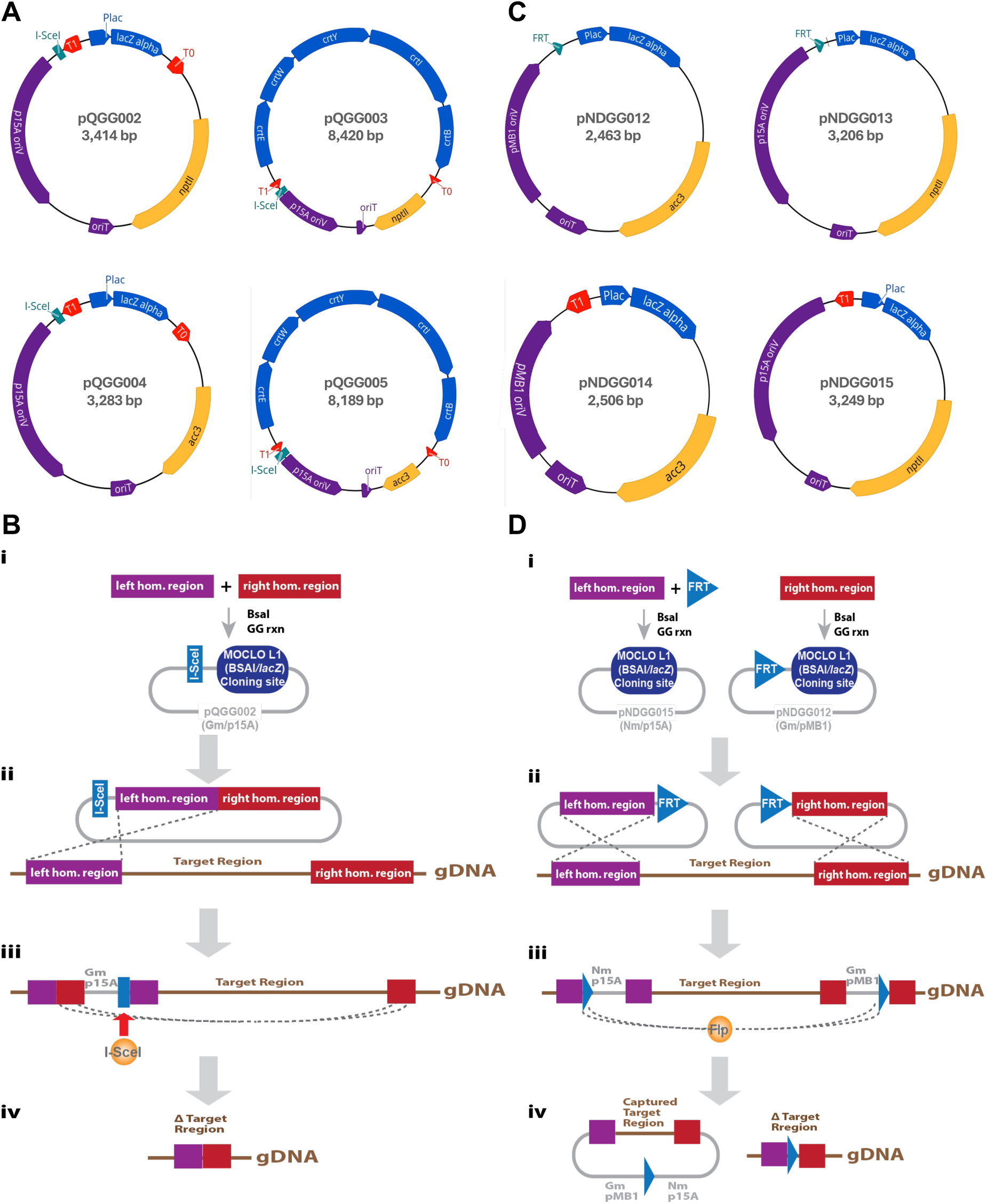
BEVA2.0 Genome Manipulation Plasmids. **A.** Maps of plasmids developed for the double homologous recombination deletion method, and **B.** schematic of the use of the plasmids to delete a target region. Steps involve: **i)** cloning of homologous regions to the left and right of the target region into a BEVA2.0 vector using BsaI or BpiI Golden Gate cloning; **ii)** conjugating the plasmid into the target organism and selecting for integration via single cross-over homologous recombination (using antibiotic resistance on the plasmid); **iii)** introducing a replicating plasmid expressing the I*-*SceI endonuclease (eg. via pDA1-ISceI-sacB) by conjugation and selecting for its maintenance to select for the second recombination event, in which the BEVA2.0 plasmid backbone is excised from the genome together with the target region; and **iv)** screening resulting colonies for the presence of the deletion (based on PCR and loss of the resistance of the BEVA2.0 plasmid). **C.** Maps of plasmids developed for Flp/*FRT* recombination methods, and **D.** a schematic diagramming their use to excise and capture a target region. Steps involve: **i)** cloning homologous regions to the left and right of the target adjacent to *FRT* site in heterologous BEVA2.0 plasmids via BsaI Golden Gate cloning; **ii)** sequentially integrating the plasmids by conjugating them into the recipient and selecting for the appropriate antibiotic resistances; **iii)** introducing a replicating plasmid containing *flp* recombinase gene (eg. pTH2505) by conjugation and inducing Flp expression to catalyze excision of the target region via *FRT* recombination (optionally also introducing an *E. coli* recipient strain for region capture, and selecting with an alternate media that selects for the recipient with BEVA2.0 plasmids); and **iv)** screening for the resulting deletion by PCR and loss of BEVA2.0 backbone resistance.

### Adapting the Flp/*FRT* deletion method to Golden Gate plasmid assembly

Another workflow for genetic manipulation involves the use of site-specific recombinases such as FLP, φC31, or Cre to generate deletions, inversions, or integrations (diCenzo and Finan, 2018). We have found these particularly useful for deleting or integrating large regions (Milunovic *et al*., 2014; Geddes *et al*., 2021). To adapt these workflows to Golden Gate cloning, we generated additional new integrative vectors. Since deletion workflows require sequential integration of recombinase sites flanking either side of a target, it is valuable to have a second heterologous integrative plasmid (diCenzo and Finan, 2018). It is optimal to minimize homology between the two plasmids, therefore in addition to p15A-based plasmids, we also generated integrative plasmids based on another integrative origin of replication, pMB1 and excluded the terminators that flank the cloning site in the BEVA architecture. We used an alternate Level 1 Golden Gate cloning site (vector pNDGG0022) that excluded the 3’ T_0_ terminator, and replaced the endlinker T1 terminator with an *FRT* site (vector pNDGG009). We designed a scheme wherein a *FRT* site can be incorporated into the 5’ end of a cloned homologous region from the genome by using the *FRT*-bearing vectors, or to the 3’ end of a cloned homologous region during region addition. To facilitate addition of the second *FRT* site, we generated an *FRT* site flanked by BsaI and 5’ GCTT and 3’ CGCT junctions (Figure 3C). We utilized these plasmids to construct an *idhA* deletion in *S. meliloti* RmP110 (Geddes, unpublished) and found them to function with similar efficiencies to traditional cloning plasmids (diCenzo and Finan, 2018).

### Linking the CIDAR parts archive to BEVA plasmids

The recently introduced CIDAR MoClo parts kit and CIDAR MoClo Expansion Volume I libraries (available as Kit 1000000059 and Kit 1000000161 through Addgene) include a wide variety of useful parts for bacterial engineering domesticated as defined parts for Level 1 Golden Gate cloning. Because both systems (CIDAR and BEVA) adopt MoClo (Weber *et al*., 2011), both systems adopt the same fusion sites for Level 1 cloning and, as such, the systems are generally compatible. However, individual CIDAR Level 0 parts were developed with an alternate Positional architecture than BEVA or MoClo (Weber *et al*., 2011; Iverson *et al*., 2016; Geddes, Mendoza-Suárez and Poole, 2019) (**Figure 4A-C**). In particular, while the 5’ fusion site GGAG (“A” in CIDAR) is conserved between the two systems and links to the 5’ fusion site of promoter parts, the 3’ cloning fusion site in BEVA is CGCT (“F” in CIDAR), whereas CIDAR was designed with terminators bearing a 3’ fusion site of GCTT (“E”). Therefore, to adapt BEVA to the CIDAR architecture, we recloned several CIDAR terminators to match the 3’ fusion site of BEVA plasmids (as “DF” rather than “DE” parts). Using these terminators enables researchers to utilize the wide variety of CIDAR parts for construction of open reading frames in BEVA plasmids. For example, we have cloned a variety of constitutively expressed fluorescent proteins by combining promoters, RBS, and open reading frames from CIDAR into the replicating plasmid pNDGG003 and verified their function in *S. meliloti* (**Figure 4E-H**, **Table 1**).

**Figure 4.**
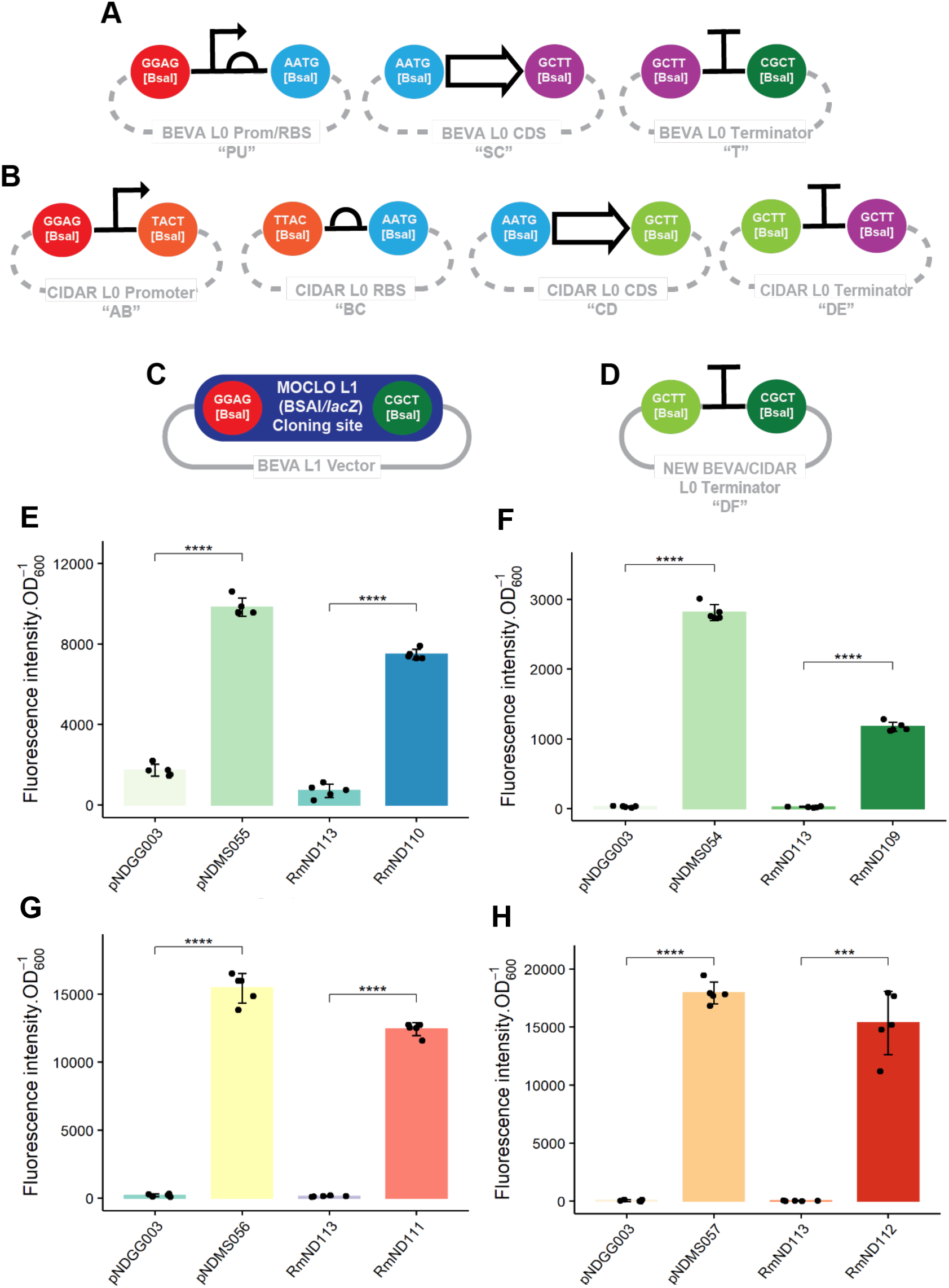
Parts to link BEVA and CIDAR. **A.** Diagram of the architecture of BEVA Level 0 parts for open reading frame construction (Geddes, Mendoza-Suárez and Poole, 2019). **B.** Diagram of the architecture of CIDAR parts for open reading frame construction. **C.** Architecture of a the BsaI multiple cloning site in Level 1 BEVA vectors. **D.** Architecture of BEVA/SEVA “linker” DF terminators. **E.** Fluorescence of BEVA vectors with CIDAR parts. The constitutive promoter J23106 together with the medium-strength RBS BCS12, are used to drive sfCFP (E), sfGFP (F), sfYFP (G) and mScarlet-I (H). The new linker terminator was used to assemble these marker genes in the RK2 BEVA2.0 plasmid pNDGG003. Fluorescence of the cloned plasmids in *E. coli* (pNDMS054-57) is shown next to the empty vector control (pNDGG003) on the left side of the graphs. On the right side of the graphs, *S. meliloti* RmP110 containing pNDGG003 (RmND113) is shown compared to RmP110 containing pNDMS054-57 (RmND109-112).

## Conclusions

The original Bacterial Expression Vector Archive has facilitated new synthetic biology approaches for alpha-proteobacteria such as advanced strain barcoding (Mendoza-Suárez *et al*., 2020) and adaption of CRISPR base editing (Wang *et al*., 2021), and has supported a range of studies by streamlining genetic engineering (Geddes *et al*., 2019; Haskett *et al*., 2022, 2023). In addition, adaptations of the BEVA architecture have recently been performed for the development of a suite of plasmids for Tn*7* (Jorrin *et al*., 2024) and mariner transposition (Williamson et al. in Preparation). The resources presented here in the BEVA2.0 expansion will further expand the utility of the system by extending the host range of replicating plasmids with new origins of replication and antibiotic resistance modules, introducing resources for genetic engineering by homologous or site-specific recombination, and adapting BEVA plasmids to function with the CIDAR system. Overall, BEVA and BEVA2.0 represent a significant community resource for bacterial genetics and synthetic biology.

## Acknowledgements

This work was supported by a New Innovator in Food & Agricultural Research (FFAR) grant to B. A. Geddes ID: FF-NIA21-0000000061m, an NSF Collaborative Research-PGR grant to B. A. Geddes: Award number 2243818. Research in the G.C.D. laboratory is supported by the Natural Sciences and Engineering Research Council of Canada (NSERC) through the Discovery Grants program (grant number RGPIN-2020-07000).

## Data Availability

Golden Gate cloning plasmids generated in this work and their DNA sequence information have been submitted to Addgene (Deposit 85226). Data supporting figures in the manuscript is available upon request.

